# Rapid, Reliable, and Interpretable CNV Curation Visualizations for Diagnostic Settings with SeeNV

**DOI:** 10.1101/2024.05.08.593244

**Authors:** Michael S. Bradshaw, Jishnu Raychaudhuri, Lachlan Murphy, Rebecca Barnard, Taylor Firman, Alisa Gaskell, Ryan M. Layer

## Abstract

Copy number variants (CNVs), structural alterations in the genome involving duplication or deletion of DNA segments, are implicated in various health conditions. Despite their clinical significance, accurate identification and interpretation of CNVs remain challenging, especially in the context of whole exome sequencing (WES), which is commonly used in clinical diagnostic laboratories. While WES offers economic advantages over whole genome sequencing (WGS), it struggles with CNV detection due to technical noise introduced by laboratory and analytic processes. Manual curation of CNV calls generated by these tools is labor-intensive and error-prone. To address this, we introduce SeeNV, a command-line tool designed to aid manual curation of CNVs at scale. SeeNV is one solution to these issues developed in collaboration with and used by the Precision Diagnostics Laboratory at Children’s Hospital Colorado. SeeNV generates static infographics for each CNV, incorporating sample and cohort sequencing coverage statistics, CNV population frequency, and more, facilitating rapid and precise assessment. Using CNVs calls identified in publicly available WES and WGS samples, we show users can rapidly and reliably curate CNV calls, needing only 4.3 seconds to curate a call, achieving 0.93 precision and 0.72 recall. SeeNV is freely available for download on GitHub: https://github.com/MSBradshaw/SeeNV.

## 1. Introduction

Copy number variants (CNVs), structural alterations in the genome involving the duplication or deletion of segments of DNA, play a pivotal role in health and disease. CNVs constitute 4.8–9.5% of the human genome ^1^ and have been implicated in a spectrum of conditions including autism, schizophrenia, Crohn’s disease, rheumatoid arthritis, type 1 diabetes, obesity, and many more ^2^. The size of CNV events can vary greatly from the duplication/deletion of a few kilobases of an exon all the way to aneuoloidy. The clinical significance of CNVs underscores the need for accurate identification and interpretation in genetic diagnostic laboratories, but remains difficult because it is done with whole exome sequencing (WES) data.

WES is more widely used in clinical settings than whole genome sequencing (WGS) as it delivers higher coverage at a lower financial cost, which is crucial for making reliable calls about small rare variants ^3;4;5^. The improvements to CNV calling made by WGS and long-read sequencing are of limited utility neither have been widely adopted outside of research settings ^3^ and the adoption process may take years or decades ^6^. While WES performs well at identifying small variants like SNPs or insertion-deletions (INDELS), identifying structural variations (SV) like CNVs can be difficult. Fundamental features of WES, such as the capture step and subsequent PCR stages, cause problems with sequencing low complexity regions and dependence on GC content can lead to over or under-representation of target regions which can be easily misinterpreted as CNVs ^5^. The difficulty of this problem is evident in the substantial variability across CNV calling methods.

We found that depending on the caller there are on average 51 - 290 calls made in a human exome and that only 12-44% are real CNV events. Due to the error-prone nature of WES-based CNV calling, manual curation and inspection of the calls and their supporting data is typically required ^2^. Given a large number of calls and enrichment for false positives, in high-throughput diagnostic settings, this manual curation task can quickly become overwhelming for clinicians.

The tools most often employed for manual curation include visualizing tools such as the UCSC Genome Browser ^7^ and/or the Integrative Genomics Viewer (IGV) ^8;9^. These two tools include graphical user interfaces that are phenomenally powerful and useful when exploring a small number of samples or calls, but when assessing hundreds of calls manually uploading multiple sample-specific files and tracks, flipping through dozens of options, and zooming in or out to find an appropriate scale, is prohibitively time-consuming.

To address the technical artifacts of WES and to aid manual curation at scale, we introduce SeeNV, a command line tool that produces a static infographic for each CNV call. SeeNV includes statistics about sample and cohort sequencing coverage, CNV population frequency, and more, integrated into a single rapidly digestible figure that requires no manipulation. We found that researchers can confidently and accurately assess a SeeNV infographic in 4.3 seconds (median value) with good precision and recall (0.93 and 0.72 respectively).

The signal behind a CNV can rise from two general sources. First is biology - the signal behind real CNV events - the natural genetic variation within individuals and across populations. Second is technical artifacts -differences in the relative abundance of reads due to PCR bias, differences in library preparation or capture steps, sequencing technologies, batches of samples, or software tools used for alignment and calling, plus more. The type and effect of technical artifacts vary greatly from one lab to the next thus technical artifacts must be controlled for by modeling lab-specific noise and signal. SeeNV does exactly this - it controls for the natural and technical sources of variation specific to one lab by relying on a reference database of samples. The first step in using SeeNV is to create a reference database of samples all processed in as exactly the same manner as possible. This set of reference samples then allows SeeNV to account for both natural biological signal and that of lab-specific technical noise at the same time.

SeeNV was originally developed for internal use at Children’s Hospital of Colorado (CHCO) and has been in their clinical pipeline for several years, playing an instrumental role in accessing the analytic reliability of CNV calls. A generalized clinical genetics workflow is shown in Figure 1, highlighting where SeeNV and it’s plots are used by CHCO. SeeNV is included as part of CHCO’s automated clinical WES pipeline, automatically generating plots for all CNV calls. The resulting infographics are then used during technical review of variants to determine if a CNV is real and worth sending on for scoring based on the American College of Medical Genetics (ACMG) criteria and clinical interpretation. We highlight two vignettes of patients at CHCO where a CNV curated with SeeNV was the diagnostic event.

**FIGURE 1.**
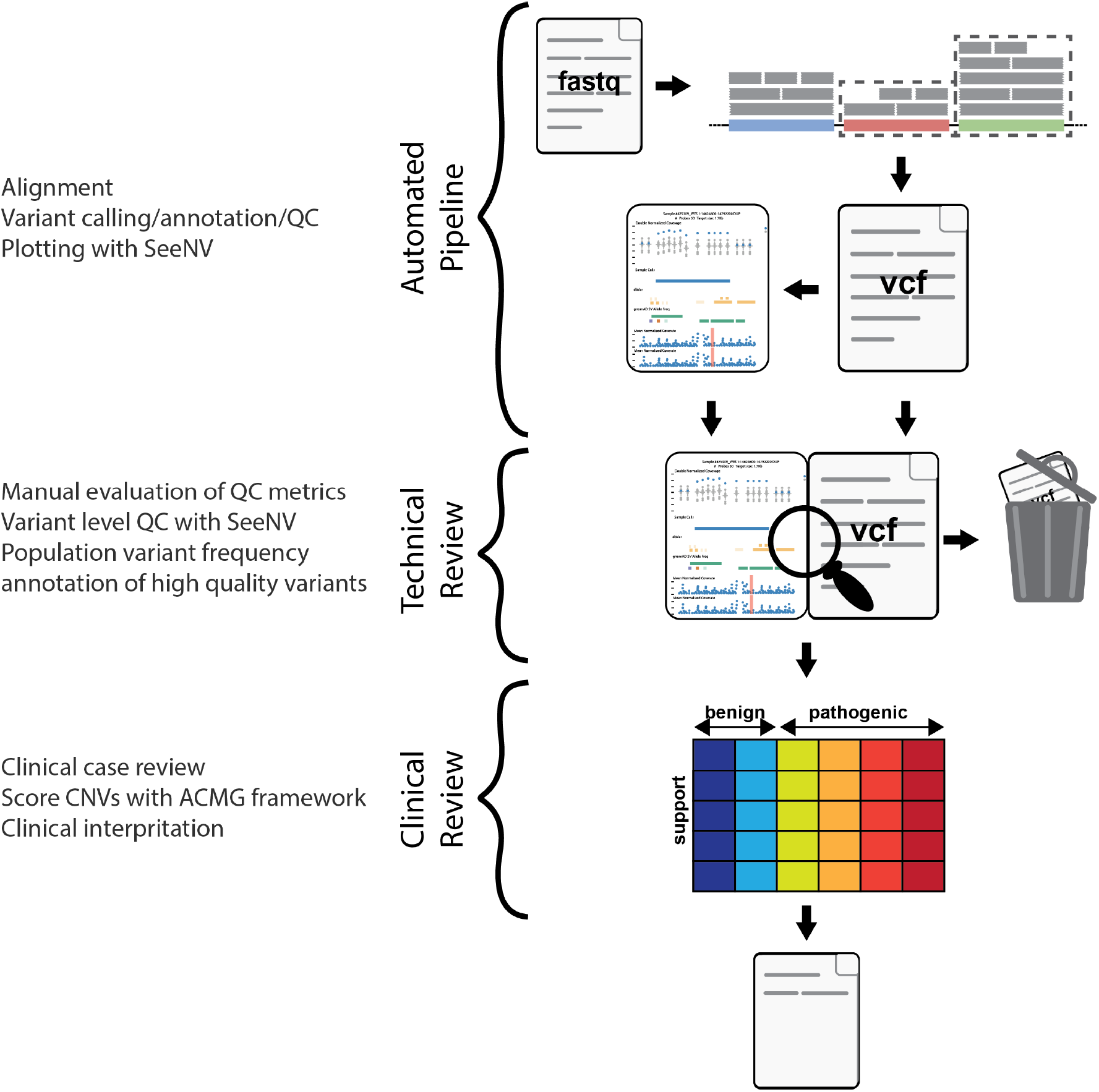
An example clinical workflow is shown divided into pipeline, technical and clinical review portions. **Automated pipeline**: a typical WES pipeline will align reads, call variants/CNVs, annotate the VCF, and perform quality control (QC). Following those steps the CNV calls and aligned reads can be fed into SeeNV for visualization. **Technical review**: final VCFs, QC reports and SeeNV infographics are used to assess the technical validity of calls. Calls determined to be real are then annotated with population allele frequency. **Clinical review**: relevant CNVs are then scored according to the ACMG framework followed by clinical interpretation and reporting of variants.

To make SeeNV readily usable by others we provide a reference database made up of 150 samples for new sam- ples to be compared and plotted against. SeeNV has built-in parallelization, a simple download and single-step install process, and an accompanying conda environment to manage dependencies. It is an open-source project available for download and use on GitHub at https://github.com/MSBradshaw/SeeNV.

## 2 Results

Here we highlight two vignettes from CHCO where a CNV identified with the help of SeeNV infographics was the diagnostic event for two patients (Figures 2 and 3). We also present the results of a user experiment where researchers were task curating WES CNV calls visualized with SeeNV and report their performance by comparing their decisions to the ground truth about these CNVs confirmed by WGS.

**FIGURE 2.**
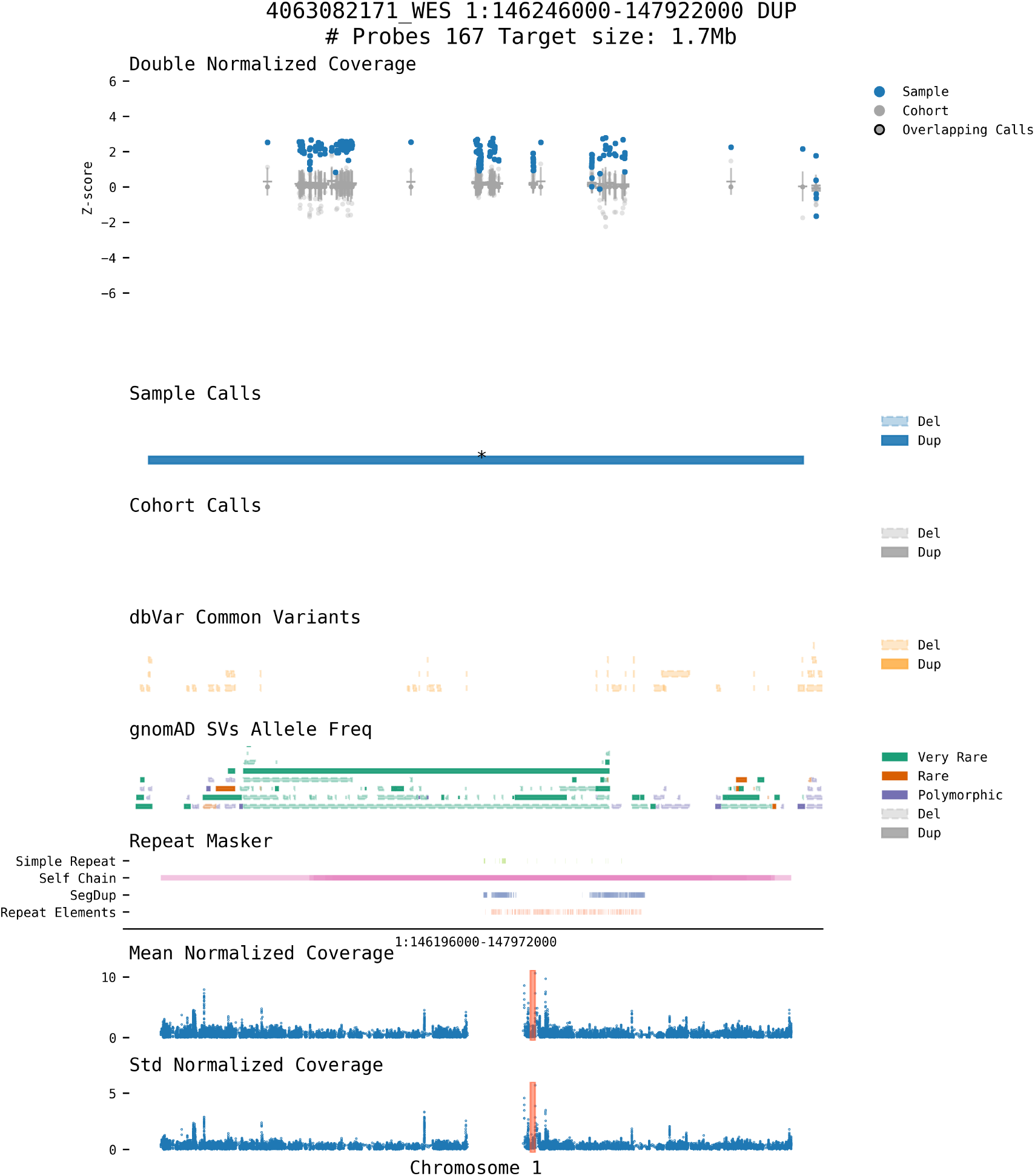
SeeNV infographic of a CNV duplication identified as the diagnostic events for a CHCO patient.

**FIGURE 3.**
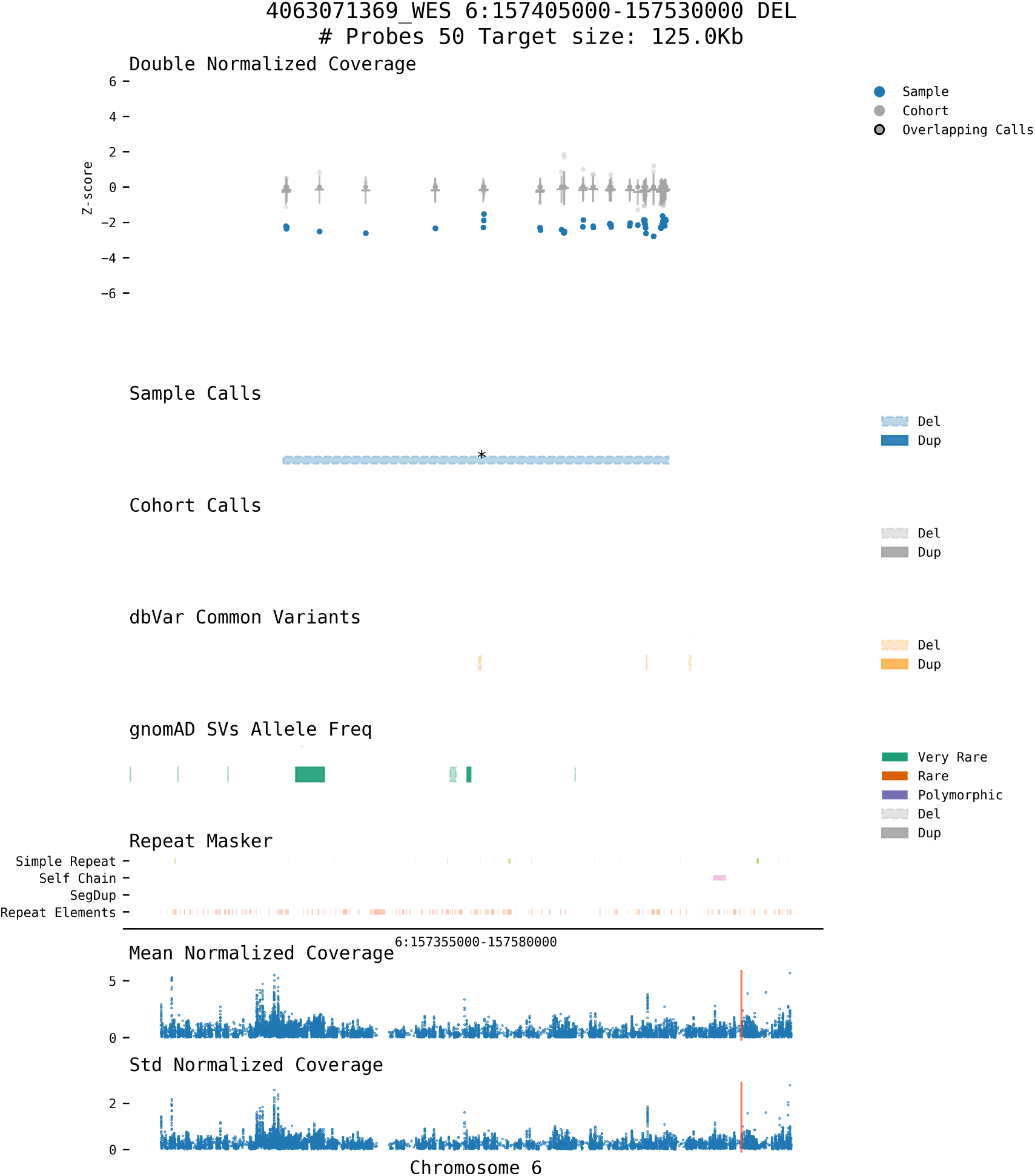
SeeNV infographic of a CNV deletion identified as the diagnostic events for a CHCO patient.

### 2.1 CHCO

Highlighted in (Figure 2 & 3) are two CNV calls from two CHCO patients where SeeNV was instrumental in helping assess the quality of a CNV that was the diagnostic event in each patient’s case. Figure 2 shows a large 1.5Mb duplication, a strong positive signal can be seen in z-scores of the call region of this sample relative to the samples in reference DB and cohort. This CNV overlaps with 173 exons from 11 genes. The patient presented the following phenotypes described with with the HPO terms: autism, abnormality of the philtrum; delayed speech and language development; sacral dimples, generalized hypotonia, obesity, pes planus, delayed gross motor development, clinodactyly of the 5th finger. A deletion is shown in Figure 3, note the negative coverage z-score of the sample at all probes overlapping with the call. This 125Kb deletion overlaps with 17 exons of the gene *ARID1B*. The patient presented the following phenotypes described with human phenotype ontology (HPO) terms: autism, short attention span, seizure, dysarthria, global developmental delay, and abnormal movement.

### 2.2 User experiment with 1000 Genomes Data

We evaluated researcher’s ability to make accurate curation decision with SeeNV using a set of CNV called in 300 WES 1000 Genomes Project Samples by the tools SavvyCNV, CNVkit and GATK gCNV. Using the WGS-validated CNVs we set up an experiment to see how reliably a researcher can curate WES CNVs using SeeNV infographics. We found that with this minimal instruction, these researchers correctly curated the vast majority of real CNVs and a majority of the false CNVs too, achieving a mean precision of 0.93 and recall of 0.72. (Figure 4.A). Mean response time of 7.5 seconds, median 4.3 seconds.

**FIGURE 4.**
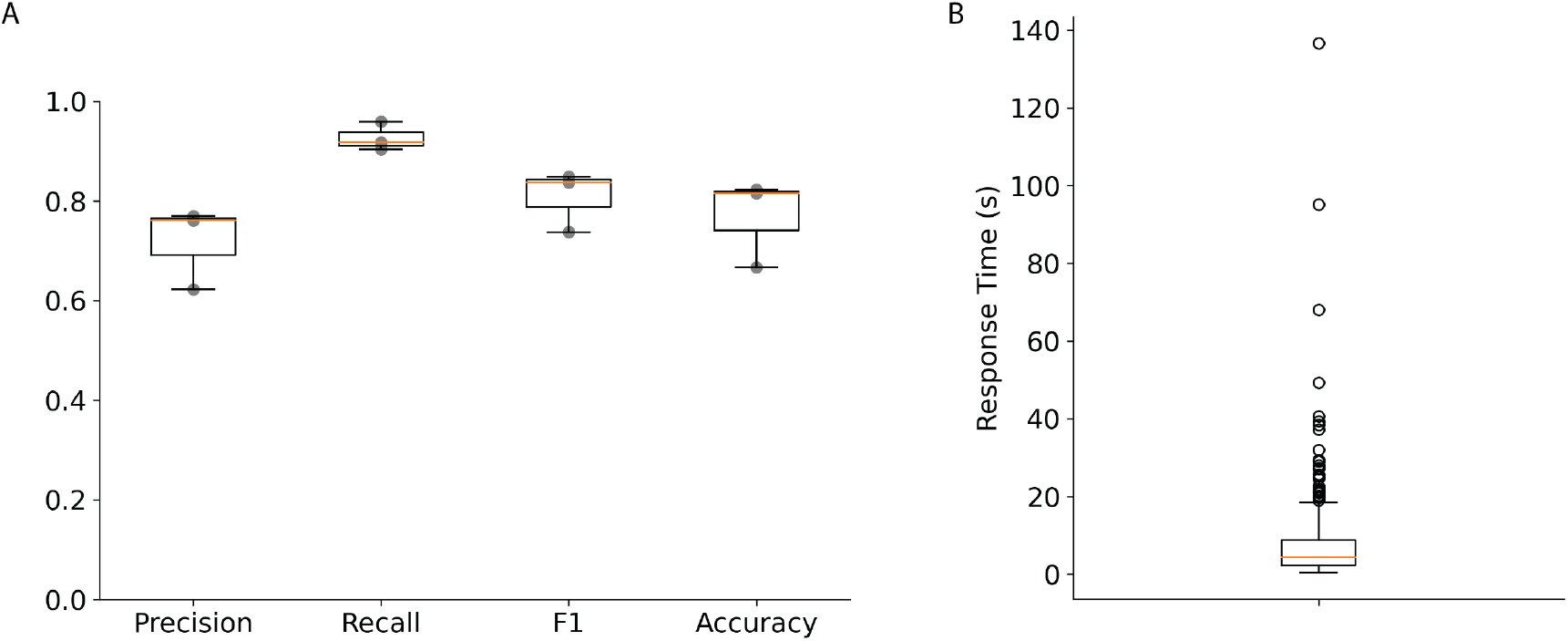
A. Results of SeeNV CNV curation experiment. Researchers were asked to label 140 CNVs visualized with SeeNV as real or not real CNVs. The truth set for evaluating the curation results was created by finding which CNVs were and were not validated by WGS SVs. B. Distribution of response times for each call.

## 3 Methods

The methods we used and document in this section include the selection of samples from CHCO and the 1000 Genomes Project, the act of CNV calling using multiple callers, establishing criteria for determining if two calls are the same, the establishment of a gold standard set of CNVs, the user curation experiment and an explanation of the parts of and statistics behind the SeeNV infographics.

### 3.1 Children’s Hospital Colorado Samples

Working with the Precision Diagnostics Laboratory at Children’s Hospital Colorado (CHCO) we called CNVs in the WES data from 721 of their patients. Permission to use this data was granted by CHCO, the study and protocol were approved by the Colorado Multi-institution Internal Review Board, ID APP001-1. All samples were deidentified prior to use and processed using the IDT xGen Exome Hyb Panel. We used three different CNV calling tools, GATK gCNA ^10^, Savvy CNV ^11^, and CNVkit ^12^.

### 3.2 1000 Genomes Project Samples

We performed the same analyses using 300 samples from 1000 Genomes Project as we did with CHCO data to demonstrate that SeeNV generalizes and to provide a reference DB for users and usage examples. Because WES is highly affected by the capture technology ^13^ and each center participating in the 1000 Genomes Project used a different capture kit for phase 3 of the project ^14^ https://www.internationalgenome.org/category/data-access/, we were limited to using samples all processed at the same center. Since so many genomic variants are ancestry-specific, databases comprised entirely of individuals from the same ancestral population can reduce or alter our ability to recognize true variation ^15^, to avoid the issues caused by this lack of representation and diversity we selected a group of samples that was perfectly balanced by ancestral population and sex. We found that choosing samples processed at the Broad Institute using Agilent SureSelect All Exon V2 capture kits provided us with the greatest number of samples while remaining balanced across sex and ancestral populations. We were able to choose 6 samples of each sex from 25 populations for a total of 300 samples, we call this set of samples the BI 300.

### 3.3 Calling Copy Number Variations

There are dozens of CNV calling tools relying on a wide variety of underlying algorithms ^1^; we use three different tools, GATK gCNA ^10^, Savvy CNV ^11^, and CNVkit ^12^. Using these three callers, we made calls in the CHCO and BI 300 samples. No effort was made to filter or quality control the output calls, we accept the output of these callers at face value. Since these callers require a reference population of samples, both sets of samples were randomly divided in half to create a set to serve as the reference and another set to use in the analyses.

### 3.4 Caller curation user experiment

Using the calls from the BI 300 and the ground truth set from WGS CNVs, we randomly selected 140 WES CNV calls to use as a test set. The test set included equal parts validated and invalidated calls as well as equal parts deletions and duplications. Infographics were loaded into PlotCritic ^16^ for ease of use in recording researchers’ curation decisions. Researchers were asked to label the call in each infographic as “yes” - a real CNV based on data in the SeeNV plot or “no” - not a CNV.

A brief explaination of the parts of the infographics was given to researchers, in addition to the text descriptions of the sections found in Section 3.5.1, and a gallery of 342 SeeNV infographics of false CNV calls and 268 true CNVs events to familiarize themselves with. In the results section we report the precision, recall, F1 and accuracy of these researchers curation decisions efforts.

### 3.5 SeeNV technical details

SeeNV is contained within its accompanying conda environment to manage dependencies. The back-end of the tool is a series of snakemake pipelines; unified modeling language (UML) diagrams of exactly how the pipelines are structured can be found in the supplemental material (Supplemental Figures 5 and 6). Currently, SeeNV has only been tested in Linux operating systems. To use and install SeeNV the only requirement is that users have conda installed in their path. There are two functions in SeeNV: build (to build a reference database (DB)), and plot (to plot CNVs). Full documentation of the parameters, usage, examples of all input files, and a small demonstration dataset can be found on the tool’s GitHub page: https://github.com/MSBradshaw/SeeNV.

#### 3.5.1 SeeNV plot explained

An example SeeNV visual from a call identified in a CHCO patient is shown in Figure 5. The information shown in the infographic from the top down is as follows:

**FIGURE 5.**
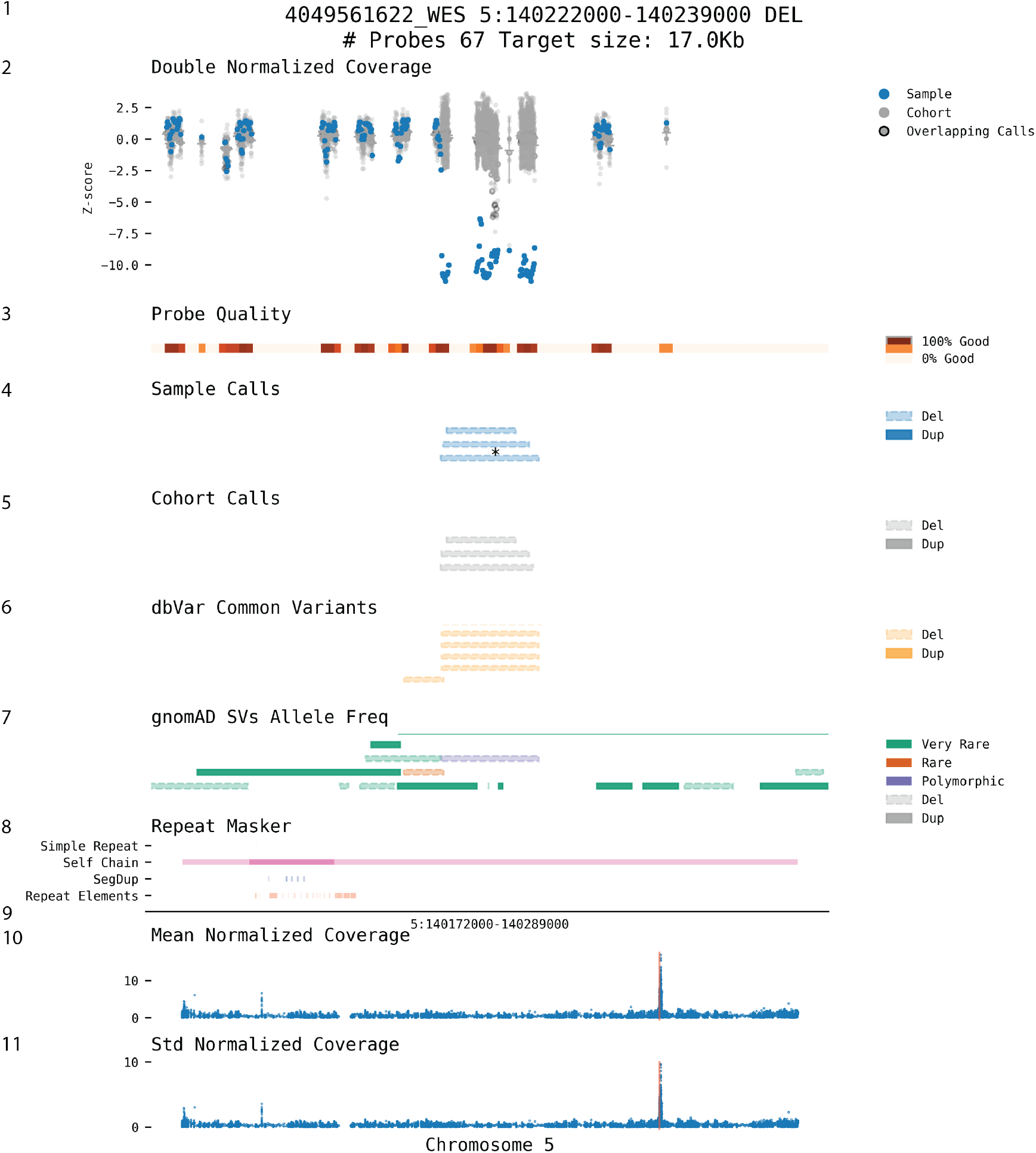
SeeNV infographic of a deletion, section numbers have been added to correspond with their in-text descriptions.

1. Title contains the sample name provided in the sample list file, genomic coordinates for the call being shown, CNV type, the number of probes the call overlaps, and the size of the window being displayed (call size + padding on either side).
2. Double normalized coverage shows the relative amount of coverage at each probe in the window. The blue dots are the in-sample probe coverage and grey dots are probe coverage from the samples in the reference DB and cohort. Grey dots with black outlines are reference DB or cohort samples that also have a call overlapping with the shown window. The horizontal lines mark the mean coverage of each probe and the vertical lines mark one standard deviation from the mean. The y-axis is a z-score of normalized coverage. Normalized coverage is calculated as:

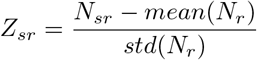

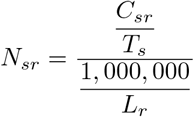

Where *N*_*sr*_ is the normalized coverage for region *r* in sample *s, Z*_*sr*_ is the z-score of the normalized coverage for region *r* in sample *s. C*_*sr*_ is the coverage or number of reads aligning to region *r* in sample *s* as identified with mosdepth mpileup. *T*_*s*_ is the total number of reads in sample *s* and *L*_*r*_ in the length of regions *r* in base pairs.
3. Probe quality is an optional section of the plot. In the bed file containing locations targeted by the specific capture technology used for the sample, you can add a column that labels certain probes as “Good” quality based on your chosen quality control metrics. This section of the plot then divides the window into 100 bins and calculates the percentage of probes labeled “Good” in each bin.
4. Sample calls shows the location of all in-sample calls falling into the window. The specific sample being plotted is marked with an asterisk. Duplications are marked with dark blue with solid edges lines and deletions are marked with a lighter blue with dashed edges. Calls that extend beyond the range of the window have the ends marked with double arrows.
5. Cohort calls shows the location of all out-of-sample calls (reference DB and cohort). Duplications are marked with dark grey with solid edges lines and deletions are marked with a lighter grey with dashed edges. Calls that extend beyond the range of the window have the ends marked with double arrows.
6. dbVar common variants marks the locations of all known common variants from dbVar. Duplications are in darker yellow, and deletions in lighter yellow with dashed edges.
7. gnomAD SVs allele freq shows the locations of SVs from gnomAD colored by their allele frequency (AF) and type. Very rare (*AF ≤* 0.01%) - green, rare (0.01% *≤ AF ≤* 0.05%) - burnt orange, polymorphic (*AF ≥* .05%) - purple. Duplications are darker shades and deletions are lighter with dashed edges.
8. Repeat Masker shows the location of various types of repeats (simple repeats, self chains, segmental duplications (SegDups), and repeat elements) collected from Repeat Masker.
9. X-axis - all plots from double normalized coverage down to Repeat Masker share this same x-axis, a window around the call in question. All plots below this x-axis are shown at chromosome scale.
10. Mean normalized coverage shows the mean normalized coverage (*mean*(*N*_*r*_)) of all samples at each region across the whole chromosome. The vertical red rectangle highlights the window being shown in the plots in the upper portion of the infographic. Typically there is one fairly large space corresponding with the centromere of the chromosome.
11. Std normalized coverage shows the standard deviation of the normalized coverage (*std*(*N*_*r*_)) of all samples at each region across the whole chromosome.

## 4. Discussion

Identifying CNVs is a difficult task that has been continually evolving. Due to its advantages in terms of economics and identifying small coding variants, WES remains more common than WGS in clinical settings.

The outstanding issues in this field we addressed here with SeeNV were the uncertainty involved in CNV calling, controlling for natural and technical artifacts affecting the coverage of targeting sequencing approaches, calling tools producing different call sets when given the same input data, and the labor-intensive task of manually curating these calls that tend to be enriched with false positives. Our research addresses these challenges head-on. Typically the statistics and visuals about a CNV used in a curation decision require multiple tools and extensive user interaction to view. SeeNV alleviates this problem by programmatically creating ready-to-view infographics that are rapidly digestible by researchers. We found that using SeeNV researchers were capable of correctly curating nearly all true CNV events (93% recall) with remarkable speed needing a median of only 4.3 seconds to curate a CNV with SeeNV. Given that WES-based CNV callers seldom agree (0.08% all calls identified in the 1000 Genomes Project samples were identified by all three callers) and output call sets are enriched with false positives (70% CNV calls are not validated), quickly and accurately filtering CNVs is important. However, the proportion of CNV calls validated by WGS much higher when all three callers agreed (100%), reaffirming the importance of relying on an ensemble of callers and looking for their agreement. By sharing SeeNV we aim to contribute to the ongoing evolution of genomic diagnostics, ensuring a balance between diagnostic yield and the clinical interpretability of CNVs.

Our research and SeeNV are not without their limitations. The double normalized coverage z-scores in the plots scale very differently for deletions and duplications. Since this statistic is a z-score, a positive value indicates greater than average normalized coverage (large positive values typically indicate a duplication) and the opposite is true for negative values (very negative values usually indicate deletions). However, the magnitude these scores can realistically achieve is in part controlled by inherent features of the genomes. Stereotypically, each human genome will have two copies of a gene (obviously this is a gross generalization), one on each strand of the chromosome, meaning at most a gene can be deleted twice. But theoretically, there is no upper limit to the number of times that gene could be copied and duplicated. We have seen this acted out in these plots fairly often. We routinely observed z-scores greater than 10 and have even seen them as high as 25 but it is very rare to see a z-score less than -3, an important consideration when viewing these plots. The magnitude of positive values in duplications is often much greater than the magnitude of negative values in deletions.

Additionally, z-score is a parametric statistic intended for normally distributed data but the depth of coverage in sequencing experiments is not necessarily normally distributed. We tested several methods of visualizing read coverage including raw counts, Poisson, and gamma standardization. Theoretically, these other standardization methods should have been more appropriate. Practically, we found that true CNV events in difficult-to-call regions of the genome were most visually distinct with our double normalized z-scores.

Concerning the three copy number callers we used and their high false positive rate, it is well known that quality control and filtering are needed. Parameters can be tuned on these callers to improve this or post hoc steps can be implemented to filter based on call length, location, or read quality ^9^; alternatively, external standalone filtering tools can be employed ^17^. We intentionally skip this step for these analyses so we can compare callers at face value but we encourage users and researchers to use automated filtering processes to further improve the quality of their CNVs and reduce the burden of manual curation.

CNV identification from WES data is a widely used and important step in clinical settings. Sensitivity can be improved with the adoption of WGS and long read technologies, but for now CNV in WES remains a staple technique. CNV callers may not agree with each other and their calls are not without their limitations, but with SeeNV we can see them a little better.

## 6. Acknowledgments

The development of SeeNV was supported by a grant from Children’s Hospital Colorado.

## 7. Author Credit

**Michael Bradshaw**: Writing-Original draft preparation, Methodology, Software, Investigation, Writing - Review & Editing. **Jishnu Raychaudhuri**: Investigation, Validation. **Lachlan Murphy**: Investigation, Validation. **Rebecca Barnard**: Conceptualization, Data Curation. **Taylor Firman**: Conceptualization, Data Curation. **Ryan Layer**: Supervision, Conceptualization, Writing - Review & Editing. **Alisa Gaskell**: Supervision, Conceptualization, Data Curation. SeeNV: visualizing copy number variations A PREPRINT

## 8. Data and code availability

## References

[1] Gabrielaite, M. et al. A comparison of tools for Copy-Number variation detection in germline whole exome and whole genome sequencing data. Cancers 13 (2021).

[2] Zarrei, M., MacDonald, J. R., Merico, D. & Scherer, S. W. A copy number variation map of the human genome. Nat. Rev. Genet. 16, 172–183 (2015).

[3] Zhao, L., Liu, H., Yuan, X., Gao, K. & Duan, J. Comparative study of whole exome sequencing-based copy number variation detection tools. BMC Bioinformatics 21, 97 (2020).

[4] Suwinski, P. et al. Advancing personalized medicine through the application of whole exome sequencing and big data analytics. Front. Genet. 10, 49 (2019).

[5] Gordeeva, V. et al. Benchmarking germline CNV calling tools from exome sequencing data. Sci. Rep. 11, 14416 (2021).

[6] Becker, F. et al. Genetic testing and common disorders in a public health framework: how to assess relevance and possibilities. background document to the ESHG recommendations on genetic testing and common disorders. Eur. J. Hum. Genet. 19 Suppl 1, S6–44 (2011).

[7] Kent, W. J. et al. The human genome browser at UCSC. Genome Res. 12, 996–1006 (2002).

[8] Thorvaldsdóttir, H., Robinson, J. T. & Mesirov, J. P. Integrative genomics viewer (IGV): high-performance genomics data visualization and exploration. Brief. Bioinform. 14, 178–192 (2013).

[9] Koboldt, D. C. Best practices for variant calling in clinical sequencing. Genome Med. 12, 91 (2020).

[10] Babadi, M. et al. Abstract 2287: Precise common and rare germline CNV calling with GATK. Cancer Res. 78, 2287–2287 (2018).

[11] Laver, T. W. et al. SavvyCNV: Genome-wide CNV calling from off-target reads. PLoS Comput. Biol. 18, e1009940 (2022).

[12] Talevich, E., Shain, A. H., Botton, T. & Bastian, B. C. CNVkit: Genome-Wide copy number detection and visualization from targeted DNA sequencing. PLoS Comput. Biol. 12, e1004873 (2016).

[13] Rosenquist, R. et al. Clinical utility of whole-genome sequencing in precision oncology. Semin. Cancer Biol. 84, 32–39 (2022).

[14] 1000 Genomes Project Consortium et al. A global reference for human genetic variation. Nature 526, 68–74 (2015).

[15] Sirugo, G., Williams, S. M. & Tishkoff, S. A. The missing diversity in human genetic studies. Cell 177, 26–31 (2019).

[16] Belyeu, J. R. et al. SV-plaudit: A cloud-based framework for manually curating thousands of structural variants. Gigascience 7 (2018).

[17] Moreno-Cabrera, J. M. et al. CNVfilteR: an R/Bioconductor package to identify false positives produced by germline NGS CNV detection tools. Bioinformatics 37, 4227–4229 (2021).

