## Supplemental methods for "Rapid, Reliable, and Interpretable CNV Curation Visualizations for Diagnostic Settings with SeeNV"

### 1. CHCO

We found the number of calls identified by the three callers within each CHCO sample (Supplemental Figure 1.A&E), the sizes of calls and limited agreement in calls made by these callers in the CHCO samples (Supplemental Figure 1.B-D&F-H). The size and quantity difference between these three callers highlights some of the uncertainty and difficulty in CNV calling. The vast majority of calls are made by a single caller (calls are considered the same if they have 60% reciprocal overlap), several hundred calls were identified by two callers, and just a few dozen were identified by all three callers.

### 2. 1000 Genomes Data

Working with 300 samples from the 1000 Genomes Project, we overlapped calls within samples based on 60% reciprocal overlap which revealed most CNVs are only identified by their respective caller (Supplemental Figure 3 A&B). Relatively few are identified by two or more callers.

We validated the CNVs identified by Savvy, CNVKit, and gCNV using the WGS CNVs. 100% of WES CNVs identified by all three calls are validated by WGS CNVs. Outside of the trice called CNVs the portions of calls that are true-positives (those WES CNVs validated by WGS CNVs) were rather low highlighting the importance of a tool like SeeNV for curation efforts Supplemental Figure 3.C&D.

[1]Department of Computer Science, University of Colorado Boulder, Boulder, CO, USA 80309,

[2]Precision Medicine Institute, Children's Hospital Colorado, Aurora, CO, USA 80045,

*Date*: May 8, 2024.

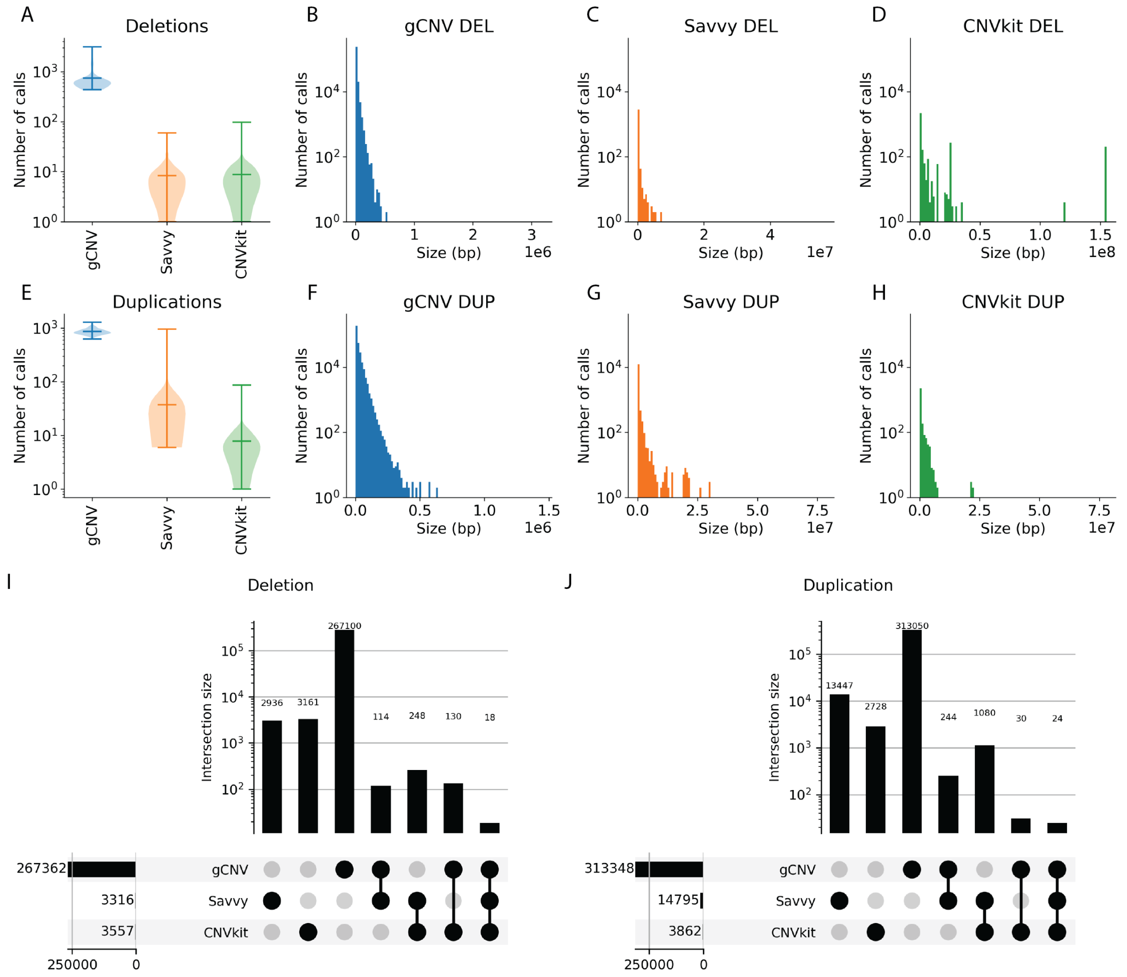


SUPPLEMENTAL FIGURE 1. The number of calls made within each CHCO sample, divided by **A**. deletions and **E**. duplications (log scale on the y-axis). Distribution of CNV sizes (y-axis on log scale) for **B**. gCNV deletions **C**. Savvy deletions **D**. CNVkit deletions **F**. gCNV duplications **G**. Savvy duplications **H**. CNVkit duplications. Upset plots showing the number of calls made in common in the CHCO samples based on 60% reciprocal overlap of **I** deletions and **J** duplication.

### 3. Determining if two CNV calls are the same

Before we could compare calls and callers we needed criteria for determining if two different calls are the same - crucial for determining if multiple samples had the same call or if two callers identified the same call in a specific sample. The sameness of calls is typically determined by intersecting the calls, however, just because two calls intersect does not mean they are the same call. Consideration needs to be made for the size of the calls and their proportional overlap. Taking this into consideration we used reciprocal overlap and an empirical approach to choosing our overlap threshold. Reciprocal overlap is when the proportion of overlap between two calls, A and B both exceed a specific threshold. In other words, for A and B to have 90% reciprocal requires that B overlaps at least 90% of A and that A also overlaps at least 90% of B.

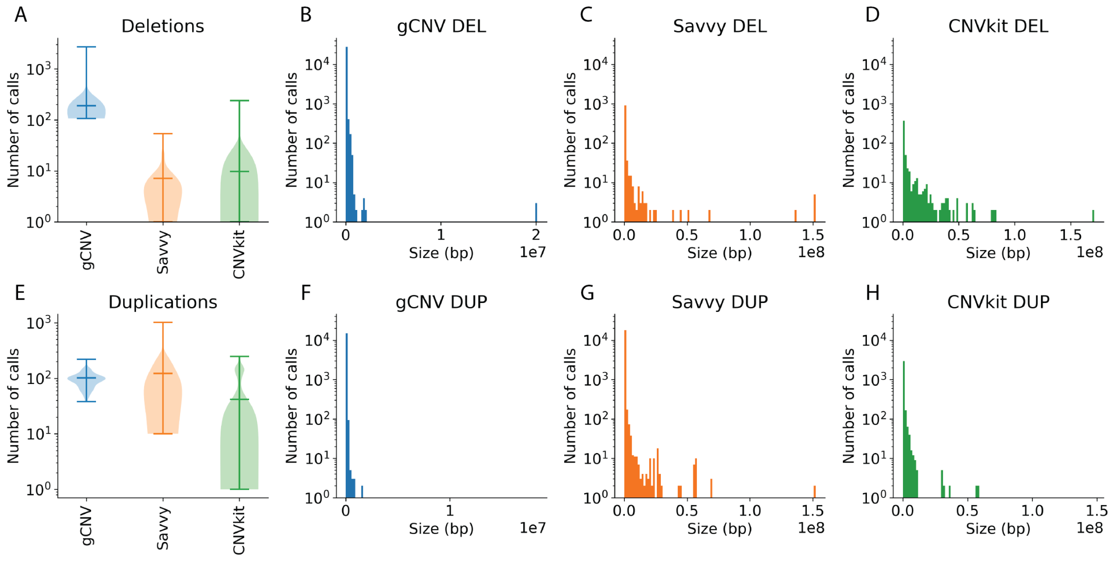


SUPPLEMENTAL FIGURE 2. The number of calls made within each BI 300 sample, divided by **A**. deletions and **E**. duplications (log scale on the y-axis). Distribution of CNV sizes (y-axis on log scale) for **B**. gCNV deletions **C**. Savvy deletions **D**. CNVkit deletions **F**. gCNV duplications **G**. Savvy duplications **H**. CNVkit duplications.

We investigated the ideal percentage of reciprocal overlap using the ‘bedtools intersect‘ function with ‘-r‘ specified and changed the value of ‘-f‘ to explore all thresholds from 0.10 to 0.95 (Supplemental Figure 4A & B). We found duplications had stable stretches of the mean overlaps per call from 0.45-0.50 and 0.55-0.65 and several steps further. The maximum number of duplication overlaps stops decreasing at 0.45, the second highest outlier stops decreasing at 0.60. For these reasons, we chose a threshold of 0.60. Deletions have continuous decay across all portions, but the maximum outlier stop decreases after .35. Due to the lack of strong evidence one way or the other for deletions, and for simplicity, we chose to use 0.60 for both types of CNVs. The cutoff of 0.60 is slightly more stringent than that used by gnomAD (0.50) for similar analyses[?] .

### 4. Determining if a CNV is validated by WGS

To create a ground truth set of real CNVs we used the CNV identified in the WGS data from the BI 300 samples in the SV supporting file from the 1000 Genomes Project download page (http://ftp.1000genomes.ebi.ac.uk/vol1/ftp/data_collections/1000G_2504_high_coverage/working/20210124.SV_Illumina_Integration/. To compare calls from the different technologies we used overlap (not reciprocal) to determine if a WGS-CNV and WES-CNV were the same. Experimenting with the overlap percentage threshold for a CNV overlapping with an SV, we determined that the ideal threshold is 0.50 for duplications and 0.30 for deletions based on the transition points in the decline of CNV to SV

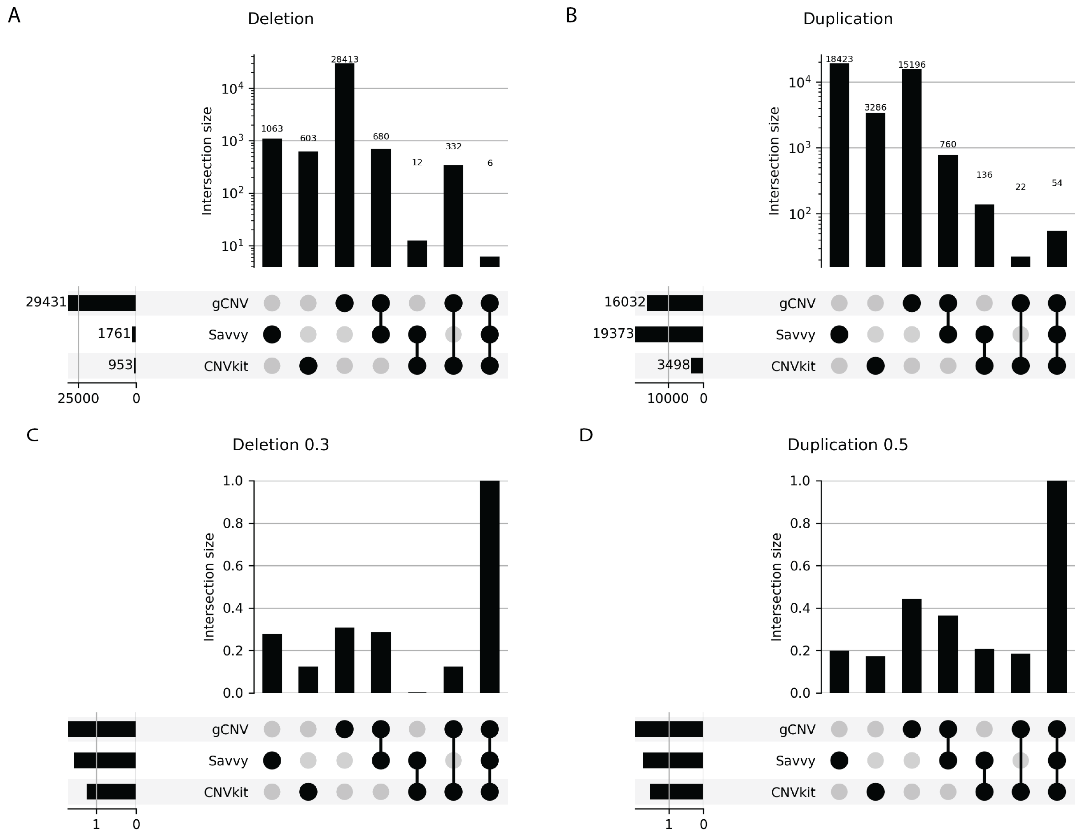


SUPPLEMENTAL FIGURE 3. Upset plots showing the number of calls made in common in the BI 300 samples based on 60% reciprocal overlap of **A** deletions and **B** duplication. Upset plots of the portion of WES-calls validated by WGS-CNVs using **C**. 30% overlap for deletions and **D**. 50% overlap for duplications.

overlaps (Figure 4C&D). These thresholds are the points where a higher threshold would make little difference in terms of the calls identified, whereas a lower one would have a large difference.

### 5. SeeNV UML Diagrams

SeeNV have two main functionalities 1 - build to create the reference database (Supplemental Figure 5) and 2 - plot to visualize CNV of input samples (Supplemental Figure 6). The workflow for these functionalities are implemented as Snakemake pipelines

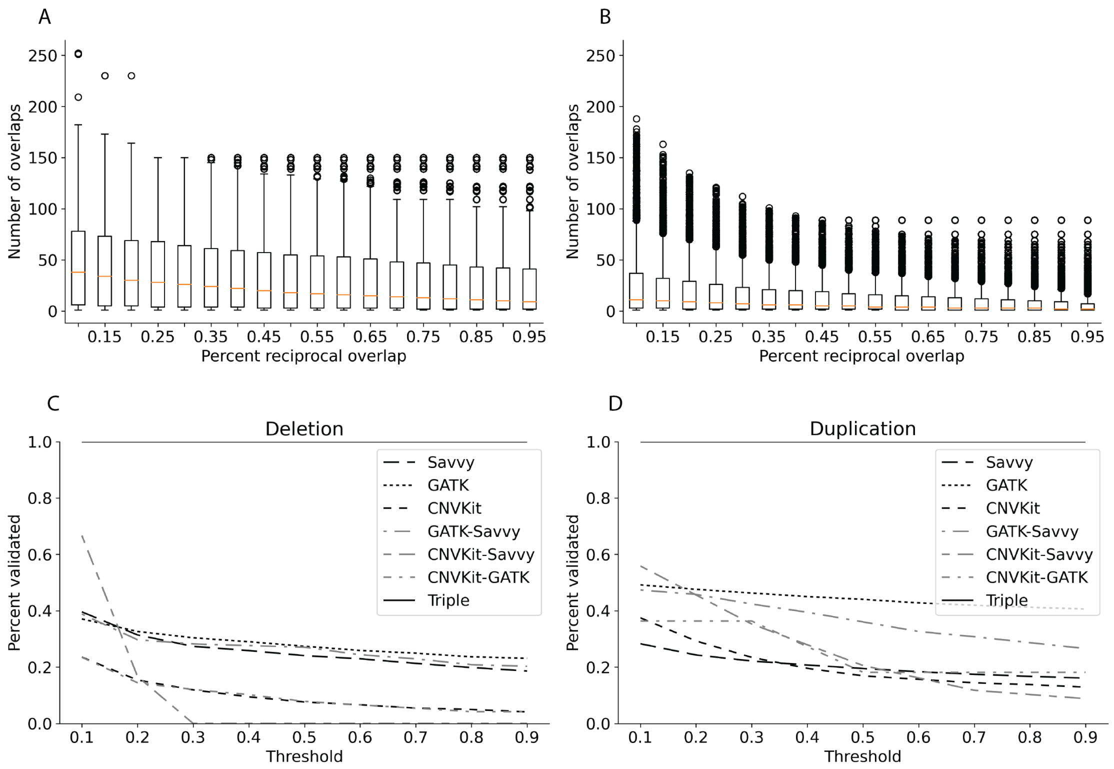


SUPPLEMENTAL FIGURE 4. The number of other calls each call overlaped with as made by gCNV, Savvy, CNVkit in the BI300 samples from 1,000 Genomes Project. **A**. The number of overlaps each deletion had with reciprocal overlap at various percent overlaps on the x-axis. **B**. The number of overlaps each duplication had with reciprocal overlap at various percent overlaps on the x-axis. The portion of calls that were validated by SVs in **C** deletions and **D** duplications, depending on the overlap threshold (x-axis).

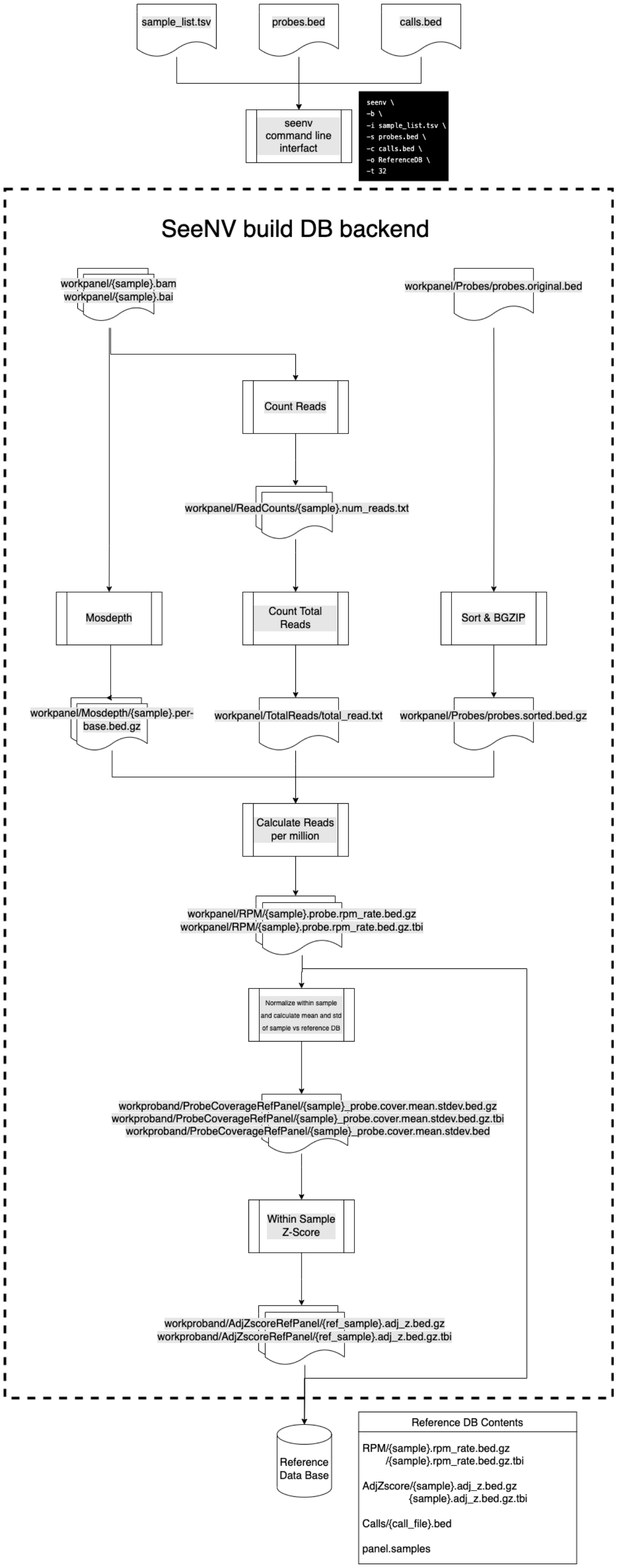


SUPPLEMENTAL FIGURE 5. Graphical view of how the build database pipeline process progresses.

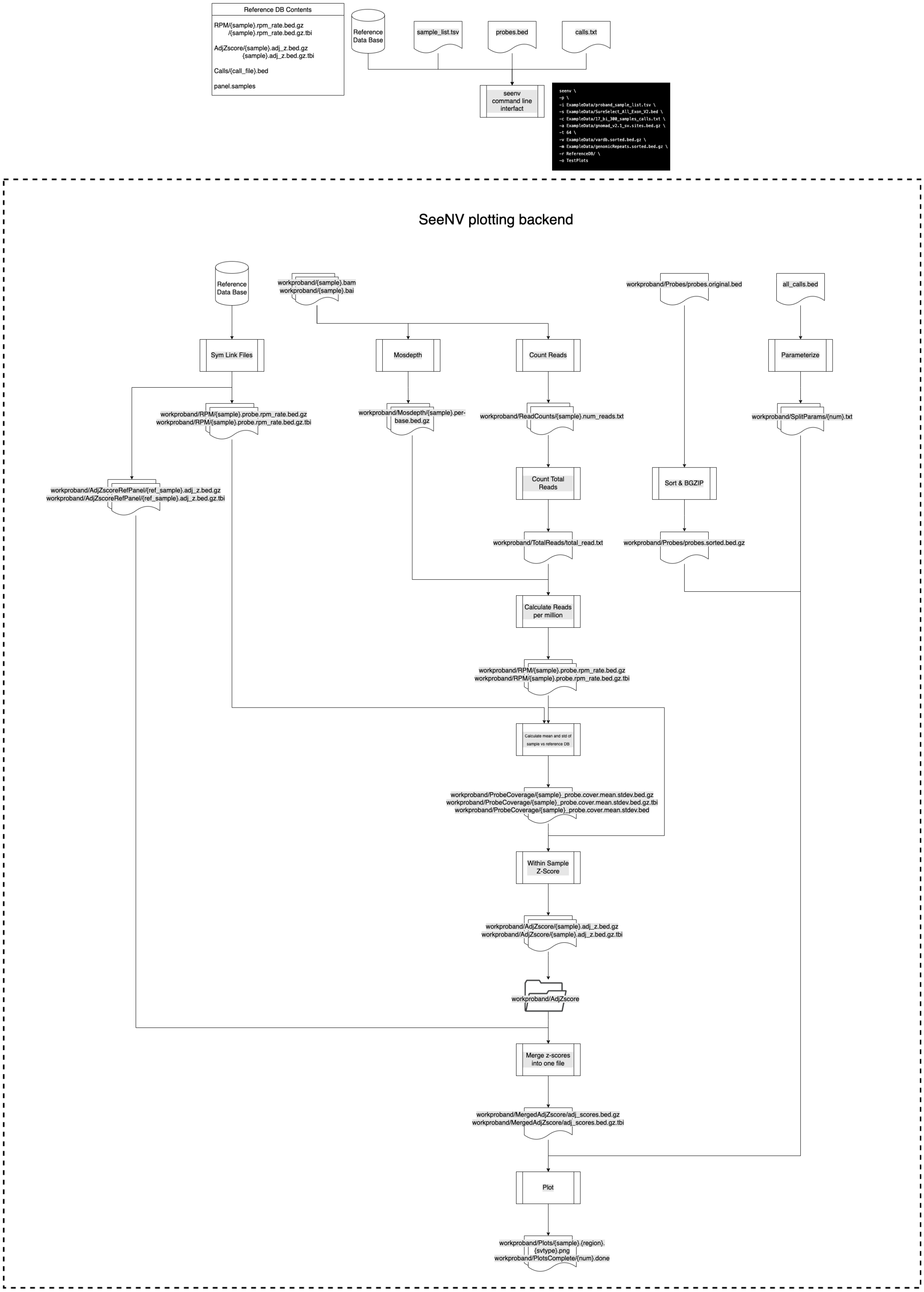


SUPPLEMENTAL FIGURE 6. Graphical view of how the plotting pipeline process progresses.
